# Gata.a, Tbx21, and Klf6/7 function cooperatively for zygotic genome activation in ascidian embryos

**DOI:** 10.64898/2026.07.30.741679

**Authors:** Kaoru S. Imai, Naoko Higuchi, Yutaka Satou

## Abstract

In ascidian embryos, gene expression from the zygotic genome begins between the 8- and 16-cell stages. While most zygotic genes are expressed in specific cell lineages at these stages, transcription factors that provide spatial cues for establishing specific expression patterns are insufficient to activate target genes at normal physiological levels. Gata.a is a transcription factor that provides spatial cues for targets expressed specifically in the animal hemisphere. Intriguingly, it is also required for physiological-level expression of many zygotic genes expressed in the vegetal hemisphere. In the present study, we found that Tbx21 and Klf6/7 augment the latter function of Gata.a. To determine the global extent of genes under control of these factors, we identified genes zygotically activated in early embryos using RNA-sequencing of BrU-labelled zygotic mRNAs. Our results revealed that approximately 80% of all zygotically activated genes were under control of these three factors. That is, together, Gata.a, Tbx21, and Klf6/7 are necessary to regulate target gene expression at physiological levels. This requirement for a specific set of broadly distributed factors resembles those of pioneer transcription factors that trigger zygotic genome activation (ZGA) in other animals, including flies and vertebrates. Regulatory factors involved in ZGA vary among animals, and our results indicate that ascidians use a distinct set of transcription factors for ZGA.

## Introduction

Maternal factors control animal development for a certain period after fertilization, during which transcription from the zygotic genome is suppressed. Thereafter, the zygotic genome begins to express genes (zygotic genome activation, ZGA). In most animals, ZGA occurs in two successive steps, the minor and major waves (Tadros and Lipshitz, 2009). In ascidian embryos, large-scale screening by *in situ* hybridization, microarrays, and sequencing has shown that zygotic gene expression begins weakly at the 8-cell stage (Ilsley et al., 2020; Imai et al., 2004; Matsuoka et al., 2013; Nishikata et al., 2001; Yamada et al., 2005). However, expression levels are much lower at the 8-cell stage than at the 16-cell stage (Oda-Ishii et al., 2018), and more genes begin to be expressed zygotically at the 16-cell stage. Thus, ZGA occurs between the 8- and 16-cell stages in ascidian embryos.

In 16-cell embryos, β-catenin, together with its effector transcription factor, Tcf7, activates genes specifically in the vegetal hemisphere, because β-catenin translocates to nuclei in the vegetal hemisphere, but not in the animal hemisphere (Hudson et al., 2013; Imai et al., 2000; Oda-Ishii et al., 2016). Meanwhile, Gata.a activates genes, including *Efna.d* and *Tfap2-r.b*, specifically in the animal hemisphere (Rothbächer et al., 2007). Although Gata.a is distributed to all cells in 16-cell embryos, its activity is suppressed by nuclear β-catenin in the vegetal hemisphere; therefore, Gata.a provides spatial cues for specific expression in the animal hemisphere (Oda-Ishii et al., 2016).

Intriguingly, Gata.a function is not limited to activating genes specific to the animal hemisphere, such as *Efna.d* and *Tfap2-r. b*. Gata.a is bound to upstream regions of genes expressed in the vegetal hemisphere, which include *Foxd*, *Fgf9/16/20*, and *Tbx6-r.b*, and is required for expression of these genes at proper levels (Imai et al., 2020). Indeed, while knockdown of *Gata.a* almost completely abolishes expression of animal-hemisphere-specific genes, it also reduces expression levels of vegetal-hemisphere-specific genes. Namely, Gata.a acts as a factor for activating animal-hemisphere-specific genes and as a factor commonly used for activating zygotic gene expression in early embryos. This latter function of Gata.a resembles those of zygotic activators. Different species use different zygotic activators, which include Zelda in *Drosophila* (Liang et al., 2008), Pha4 (*Foxa*) in *Caenorhabditis* (Hsu et al., 2015), and Pou and Sox factors in vertebrates (Gentsch et al., 2019; Lee et al., 2013; Leichsenring et al., 2013). Many of these factors exhibit pioneering activity, and Gata is also a pioneer transcription factor (Iwafuchi-Doi and Zaret, 2014).

Although Gata.a affects many genes expressed in early embryos, some genes expressed in early embryos do not contain putative Gata.a binding sites in their upstream regulatory regions (Imai et al., 2020). The extent of expression level reduction in *Gata.a* morphants also varies among genes. Several motifs other than Gata.a binding sites are also enriched in upstream regulatory regions of genes activated at the 16-cell stage (Imai et al., 2020). These observations indicate that ascidian embryos contain additional transcription factors required for activating genes at proper levels at ZGA.

## Results

### *Tbx21* and *Klf6/7* are required for proper gene expression in early embryos

We previously reported that *Tbx21*, which encodes a T-box transcription factor, is maternally expressed, and that its zygotic expression is detected ubiquitously at the 16-cell stage, except in germline cells (Imai et al., 2020). This broad expression pattern is one of the criteria for regulators of zygotic genome activation (ZGA). We indeed identified putative Tbx21 binding sites in the 1-kb upstream regions of *Efna.d* and *Tfap2-r.b* (Figure S1), two genes that initiate expression at the 16-cell stage (Imai et al., 2004). Therefore, we examined expression of *Efna.d* and *Tfap2-r.b* in embryos injected with a morpholino oligonucleotide (MO) against *Tbx21*, using *in situ* hybridization. While almost all embryos injected with a control MO, which does not have targets in the ascidian genome, expressed *Efna.d* and *Tfap2-r.b*, 82% and 96% of *Tbx21*-morphant embryos lost expression of *Efna.d* and *Tfap2-r.b* at the 16-cell stage, respectively (Figure 1A–D).

**Figure 1.**
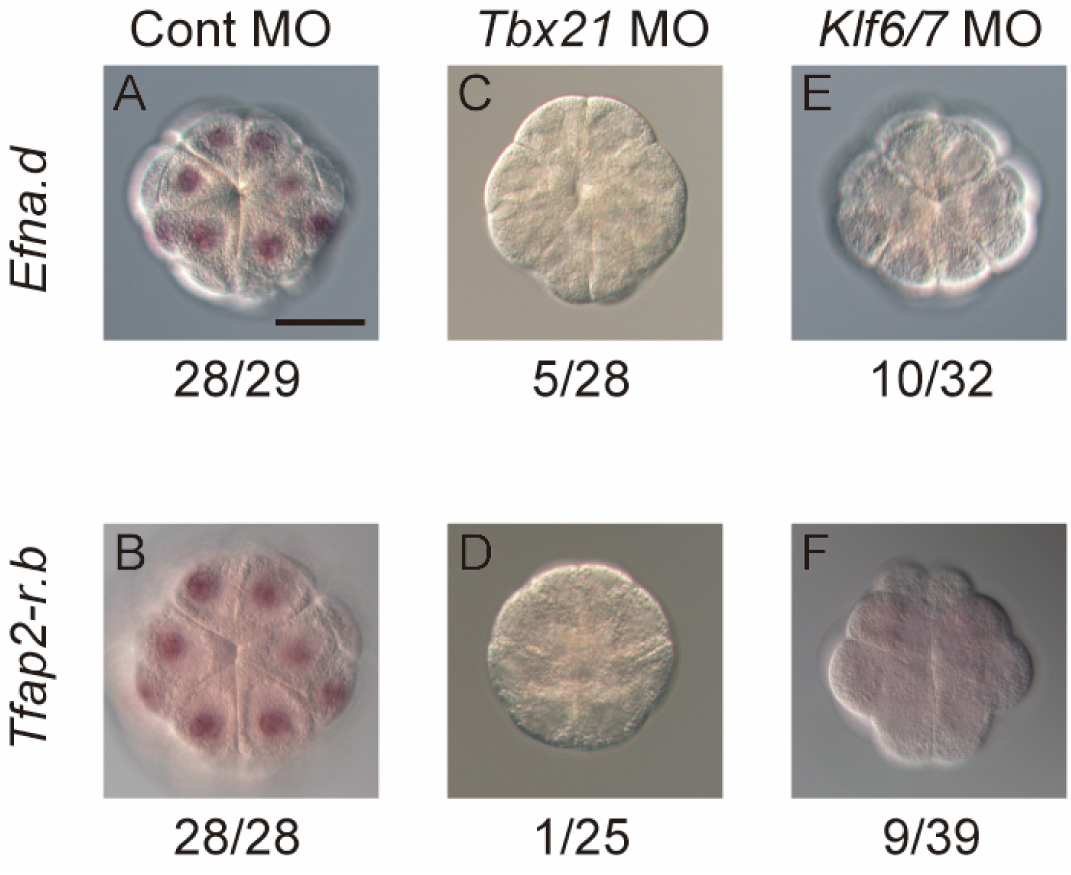
*Tbx21* and *Klf6/7* regulate expression of *Efna.d* and *Tfap2-r.b*. Expression of (A, C, E) *Efna.d* and (B, D, F) *Tfap2-r.b* in embryos injected with (A, B) the control MO, (C, D) *Tbx21* MO, or (E, F) *Klf6/7* MO was examined by *in situ* hybridization at the 16-cell stage. All experiments were performed three times with three different batches of embryos. Total numbers of embryos and numbers of embryos with expression are shown beneath the photographs. MOs were injected at 0.5 mM. Scale bar, 50 μm.

We took another approach to find additional factors that are ubiquitously required for proper gene expression. We previously identified eleven 6-mer motifs enriched in upstream regions of genes expressed in 16-cell and/or 32-cell embryos, and the top two 6-mers included ‘GATA’, which potentially bind a GATA transcription factor (Imai et al., 2020). Among the remaining enriched motifs, seven contained a ‘GCG’ triplet. Therefore, we searched the Ghost database (Satou et al., 2005) for transcription factors that preferentially recognize ‘GCG’, and mRNAs of which are expressed abundantly in eggs, but rarely in gastrula embryos. Zlf6/7 (formerly Zf219), which is an ortholog for human KLF6 and KLF7 (Tamura and Satou, 2025), met these criteria. The consensus binding sequence for Klf6/7, ‘GGGCGT’ (Figure S1B)(Nitta et al., 2019; Vincentelli et al., 2011), contains the ‘GCG’ triplet, and maternal expression has been experimentally validated (Brozovic et al., 2018; Imai et al., 2004; Matsuoka et al., 2013). Using previously published gene expression data (Ilsley et al., 2020; Matsuoka et al., 2013), we also confirmed that its mRNA is detected in all blastomeres at almost equal levels (Figure S2). Because we succeeded in identifying several putative Klf6/7-binding sites in the 1-kb upstream regions of *Efan.d* and *Tfap2-r.b*, we again examined expression of *Efna.d* and *Tfap2-r.b* in embryos injected with an MO against *Klf6/7* for *in situ* hybridization, and found that 96% and 77% of *Klf6/7-*morphant embryos lost *Efna.d*, and *Tfap2-r.b* at the 16-cell stage (Figure 1E, F).

### *Gata.a*, *Tbx21*, and *Klf6/7* regulate gene expression cooperatively in early embryos

Because *Efna.d* and *Tfap2-r.b* also require Gata.a for initiating expression (Imai et al., 2020), we examined the possibility that these three factors act cooperatively. We injected the *Gata.a* MO from our previous study (Imai et al., 2020), but at a reduced dose (0.5 mM), which was half the concentration employed previously, to ensure that cooperative effects would be more easily discernible by reverse-transcription and quantitative PCR (RT-qPCR). While expression levels of *Efna.d* and *Tfap2-r.b* in *Gata.a* morphants were approximately half of those in control embryos injected with the control MO (Figure 2A), their expression levels in double morphants (0.5 mM each) were more markedly reduced than those in any single morphants of *Gata.a*, *Tbx21* or *Klf6/7*.

**Figure 2.**
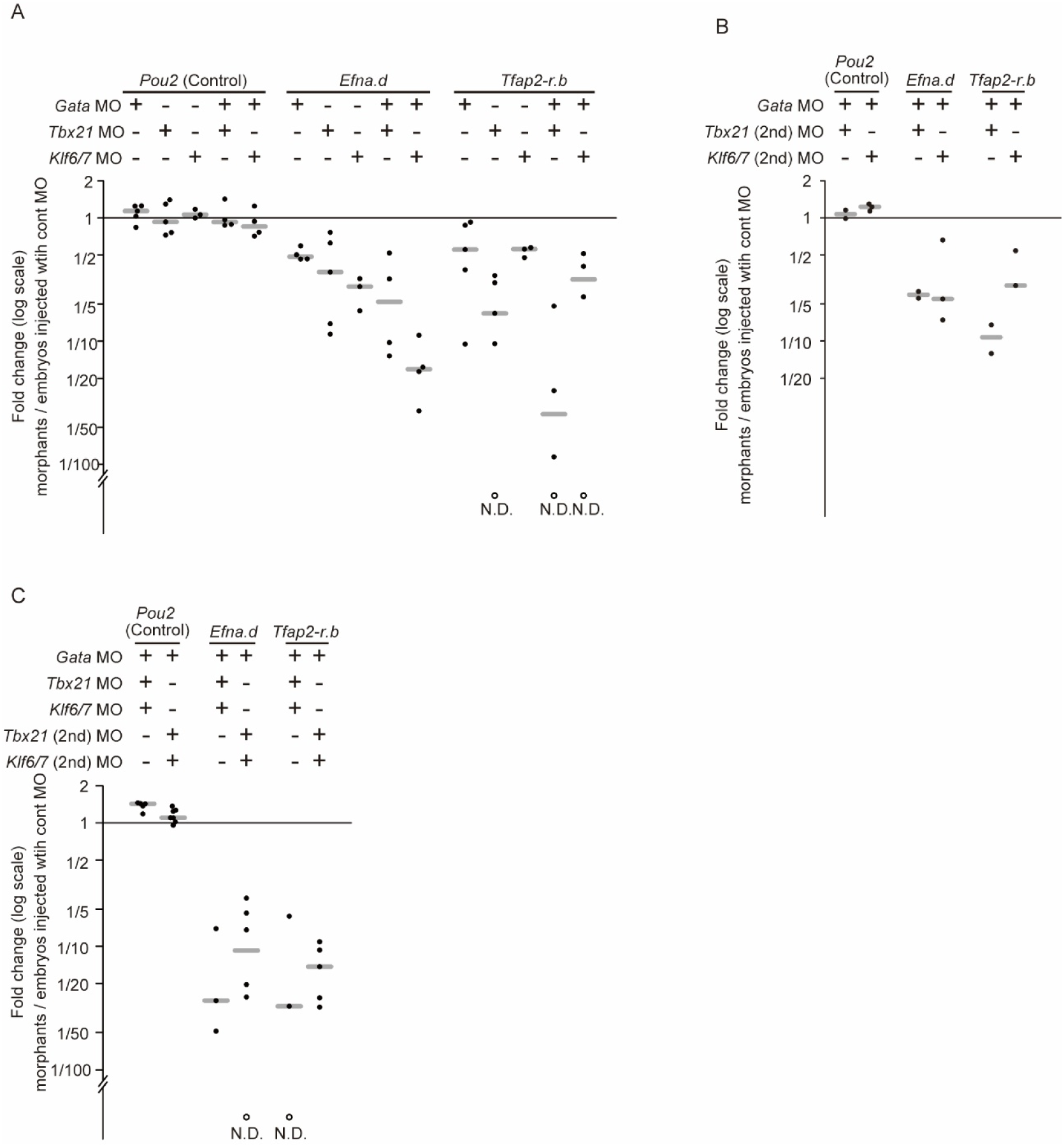
*Tbx21* and *Klf6/7* regulate expression of *Efna.d* and *Tfap2-r.b* cooperatively with *Gata.a*. Expression levels of *Efna.d* and *Tfap2-r.b* were measured by RT-qPCR. *Macho-1* was used as a reference for the ΔΔCt method. Dots represent individual experimental results (biological replicates), and gray lines indicate medians. Expression of *Pou2*, which is expressed maternally, was also examined as a control. White dots indicate that no amplification was detected. (A) Relative expression levels in 32-cell embryos injected with an MO for *Gata.a*, *Tbx21*, or *Klf6/7* (0.5 mM), or a mixture of two MOs of these three MOs (0.5 mM each) against embryos injected with a control MO (0.5 mM). (B) Relative expression levels in 32-cell embryos injected with the *Gata.a* MO and a second MO against either *Tbx21* or *Klf6/7* (0.5 mM each). (C) Relative expression levels in 32-cell embryos injected with MOs against *Gata.a*, *Tbx21*, and *Klf6/7*. Because five data points lacked values (no detection), and because similar measurements were performed in subsequent analyses, we did not perform statistical tests in these experiments.

To confirm the specificity of the *Tbx21* and *Klf6/7* MOs, we injected secondary MOs against these genes with the *Gata.a* MO. Consistent with results obtained with the primary MOs, expression levels of *Efna.d* and *Tfap2-r.b* were reduced to a greater extent in double morphants (the primary *Gata.a* MO/secondary *Tbx21* MO, or the primary *Gata.a* MO/secondary *Klf6/7* MO) than in embryos injected with the *Gata.a* MO alone (Figure 2B). These results suggest that MOs against *Tbx21* and *Klf6/7* specifically suppressed their respect targets.

Next, we injected a mixture of all three MOs against *Gata.a*, *Tbx21*, and *Klf6/7*. For this experiment, we used each MO at a concentration of 0.25 mM, which is lower than the concentrations we employed in the preceding experiments. Nevertheless, the expression levels of *Efna.d* and *Tfap2-r.b* were substantially reduced (Figure 2C).

### Identification of genes zygotically activated at the 16- and 32-cell stages

The above analyses raised the possibility that Tbx21 and Klf6/7 might be broadly required for expression of many genes in early embryos, as in the case of Gata.a. To examine the global extent to which early embryonic genes are regulated by these three factors, we prepared three types of embryos; uninjected control embryos (experiment 1), embryos injected with bromouridine (BrU) and the control MO (experiment 2), and embryos injected with BrU and a mixture of the MOs against *Gata.a*, *Tbx21*, and *Klf6/7* (experiment 3). Embryos were collected at the 16- and 32-cell stages, and RNAs were extracted from these embryos. RNAs obtained from experiments 2 and 3 were then immunoprecipitated with an anti-BrU antibody to specifically isolate zygotically transcribed mRNAs for RNA-sequencing. For each experiment, we took two specimens from different batches of embryos at each of the 16- and 32-cell stages (two biological replicates). All transcripts-per-million (TPM) are provided in Table S1.

First, we compared transcriptomes between experiments 1 and 2 to identify zygotically activated genes. We identified 4712 genes significantly enriched in specimens from experiment 2 at the 16-cell stage, and 5595 genes at the 32-cell stage (adjusted p-value < 0.001). To obtain lists of genes that are zygotically expressed at levels detectable by *in situ* hybridization, we filtered these genes as follows. Previous studies that examined gene expression by *in situ* hybridization have identified 27 genes expressed at the 16-cell stage (*Admp*, *Dusp6/7/9*, *Efna.a*, *Efna.d*, *Fgf9/16/20*, *Foxa.a*, *Foxd.a*, *Foxd.b*, *Foxtun1, Foxtun2*, *Fzd4*, *Gdf1/3-r*, *Hes.a*, *Lefty, Noto*, *Prdm1-r.a*, *Smad1/5/9*, *Sox1/2/3*, *Tbx21*, *Tbx6-r.a*, *Tbx6-r.b*, *Tbx6-r.c*, *Tbx6-r.d*, *Tfap2-r.b*, *Wnttun5, Sp.b/Zf220*, and *Zfpm*) (Ikeda et al., 2013; Imai et al., 2002a; Imai et al., 2004; Imai et al., 2002b; Takatori et al., 2010). All these genes except *Foxtun2* were present in the above list for the 16-cell stage. This confirms that Bru-RNA-sequencing successfully identified genes activated zygotically at the 16-cell stage. Among these known genes, except *Foxtun2*, *Tbx6-r.a* exhibited the lowest TPM value. Consequently, we used its TPM as a threshold, and identified 383 genes that had TPM values equal to or exceeding this threshold (Figure 3A). We regarded this as the set of genes zygotically expressed at the 16-cell stage (Table S2).

**Figure 3.**
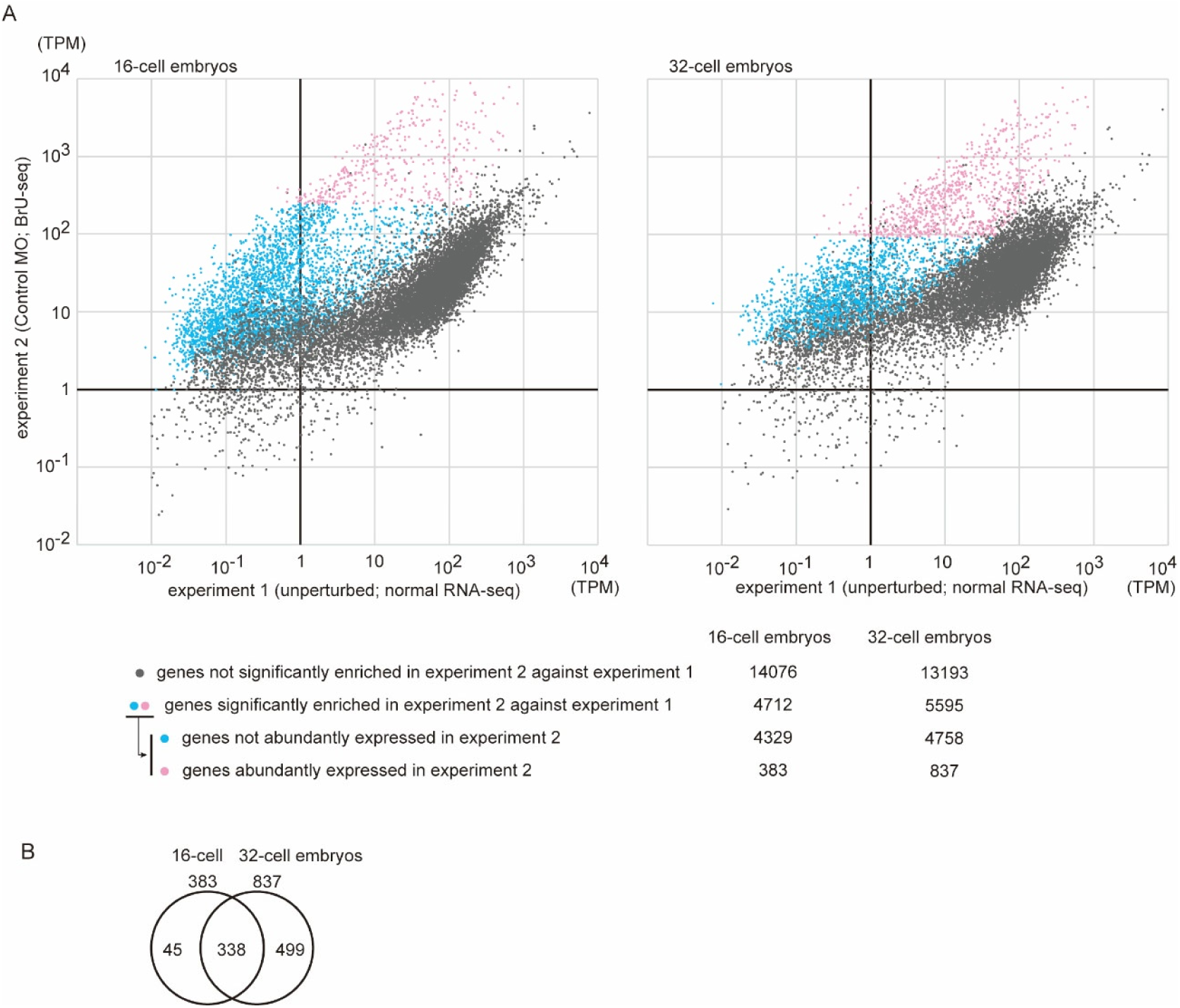
Genes zygotically activated in 16- and 32-cell embryos were revealed by Bru-RNA-sequencing. (A) RNA-sequencing reads were mapped to the HT version of the genome (Satou et al., 2019) and the KY21 gene model sets (Satou et al., 2022), and transcripts-per-million (TPM) were calculated. Genes were plotted according to their TPM values obtained from experiments 1 and 2. Genes were categorized into three groups and represented by three colors. Meanings of colors are shown below the graphs with numbers of genes that individual groups contain. (B) Most genes zygotically expressed at the 16-cell stage are also expressed zygotically at the 32-cell stage.

Similarly, at the 32-cell stage, 20 additional genes are expressed zygotically (*Bmp3, Dkk, Dlx.b, Dmrt.a, Dusp1/2/4/5*, *Hes.b*, *Lhx3/4, Neurog, Nodal, Otx, Prdm1-r.b, Snai, Wnt3*, *Wnt5, Zic-r.b, Zic-r.c, Zic-r.d, Zic-r.e, Zic-r.f,* and *Zic-r.g*) (Hudson and Lemaire, 2001; Ikeda et al., 2013; Imai et al., 2004; Imai et al., 2002c; Satou et al., 2001), and all except *Foxtun2* were included in the above list for the 32-cell stage. *Dmrt.a*, which had the lowest TPM among them, was used as the threshold to filter the listed genes, and we identified 837 genes zygotically expressed at the 32-cell stage (Table S3). Of the 383 genes expressed at the 16-cell stage, 338 genes (88.3%) were included in the list at the 32-cell stage (Figure 3B).

### Many genes zygotically expressed in early embryos encode transcriptional regulators

We found that 210 of 383 genes zygotically expressed at the 16-cell stage encoded homologs of human proteins (BLASTP, E-value < 1×10^-5^). Using GO terms associated with these human homologs, we identified two molecular-function GO terms, one biological-process GO term and one cellular-component GO term that were significantly enriched among proteins encoded by genes zygotically expressed at the 16-cell stage (Figure 4). Specifically, the enriched molecular function and biological-process GO terms were related to regulation of transcription (GO:0000978, GO:0000981, and GO:0000122; Table S4). Furthermore, the cellular-component GO term was “Chromatin” (GO:0000785).

**Figure 4.**
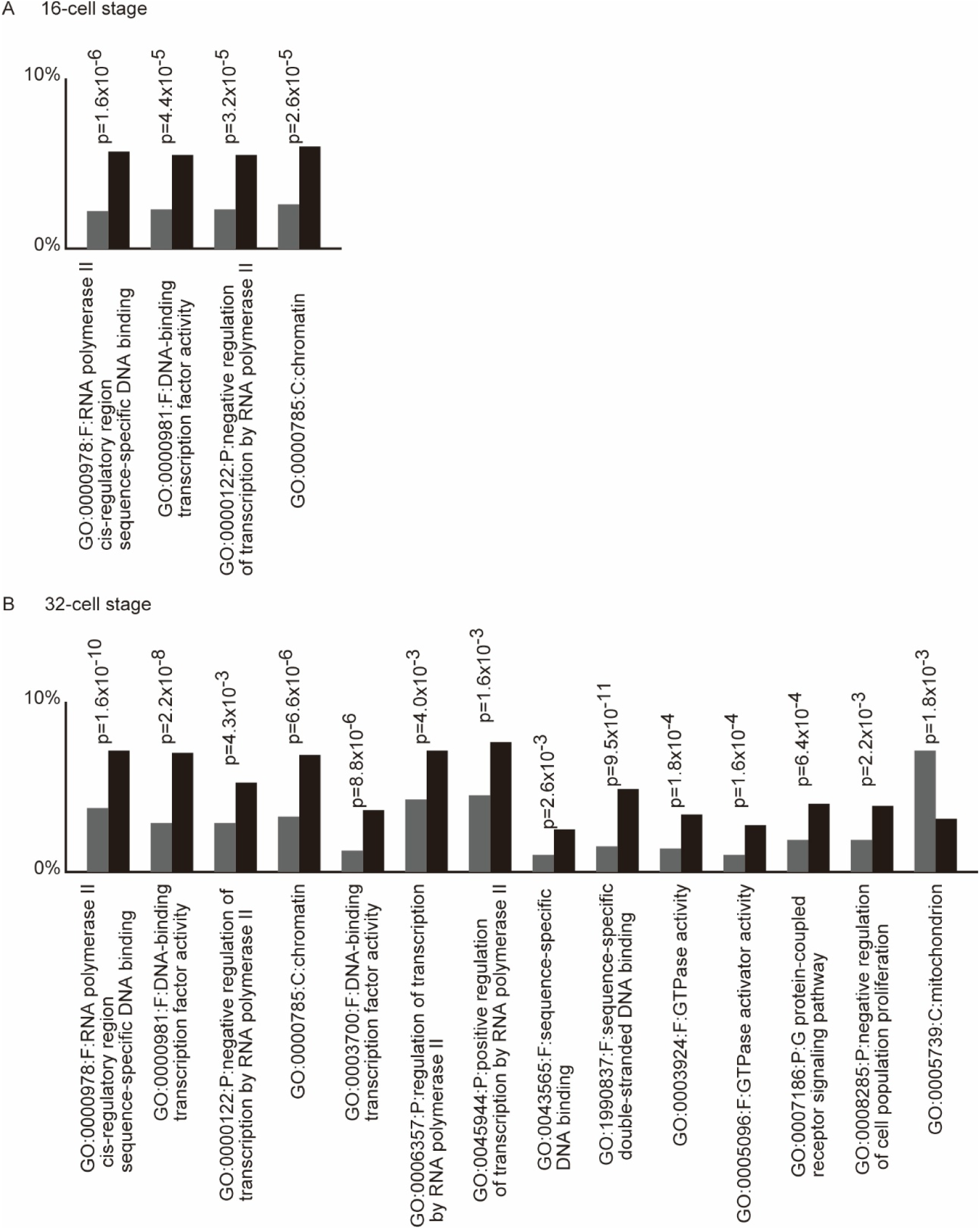
A gene ontology analysis of proteins encoded by genes zygotically activated in 16- and 32-cell embryos. Gene ontology terms over-/under-represented in products of genes zygotically activated (A) at the 16-cell stage and (B) at the 32-cell stage (light gray bars) in comparison with the entire protein set encoded in the genome (black bars). Differences were examined by Chi-square tests, and p-values are shown. Only GO terms that passed a Benjamini-Hochberg test (threshold value =0.01) are shown. Each label below the graphs includes three fields. The first field indicates the GO term number. The second indicates a GO category: F, molecular function; P, biological process; C, cellular component. The last field presents an explanation.

Similarly, among the 837 genes zygotically expressed at the 32-cell stage, 475 genes had human homologs. The four GO terms mentioned above and five additional GO terms (GO:0003700, GO:0006357, GO:0045944, GO:0043565, and GO:1990837) related to transcriptional regulation or DNA-binding were enriched at the 32-cell stage. In addition to these GO terms, four GO terms were over-represented and one was under-represented (Figure 4B; Table S5). These terms did not appear to be directly related to transcriptional regulation.

All zygotically activated genes with potential roles in transcriptional regulation are listed in Table S6. While many of them are sequence-specific transcription factors, and their expression patterns have been analyzed by *in situ* hybridization, microarrays, or RNA-sequencing (Ilsley et al., 2020; Imai et al., 2004; Imai et al., 2020; Matsuoka et al., 2013; Miwata et al., 2006; Yamada et al., 2005), we identified several genes whose zygotic transcription had not been reported. These include *Mxd*, *Foxh.b*, *Rela*, *Prdtun1, Ror, Foxp, Sox4/11/12*, and *Tcf3*. The failure of previous studies to detect these zygotic transcripts is probably due to high levels of maternal expression, which masks zygotic expression. For this reason, we were not able to confirm their zygotic expression by *in situ* hybridization; however, BrU-labeling identifies zygotically activated genes more sensitively and efficiently than methods used in previous studies.

### Gata.a, Tbx21, and/or Klf6/7 are required to activate approximately 80% of genes in 16- and 32-cell embryos

To assess the global impact of Gata.a, Tbx21, and Klf6/7, we compared transcriptomes in experiments 2 and 3. DESeq2 analysis revealed that 307 of the 383 genes (80.2%) zygotically expressed at the 16-cell stage were significantly down-regulated in triple morphant embryos injected with MOs against *Gata.a*, *Tbx21*, and *Klf6/7* (adjusted p-value < 0.001) (Figure 5A; Table S7). Similarly, 663 of the 837 genes (79.2%) zygotically expressed at the 32-cell stage were significantly down-regulated in triple morphants (Table S8).

**Figure 5.**
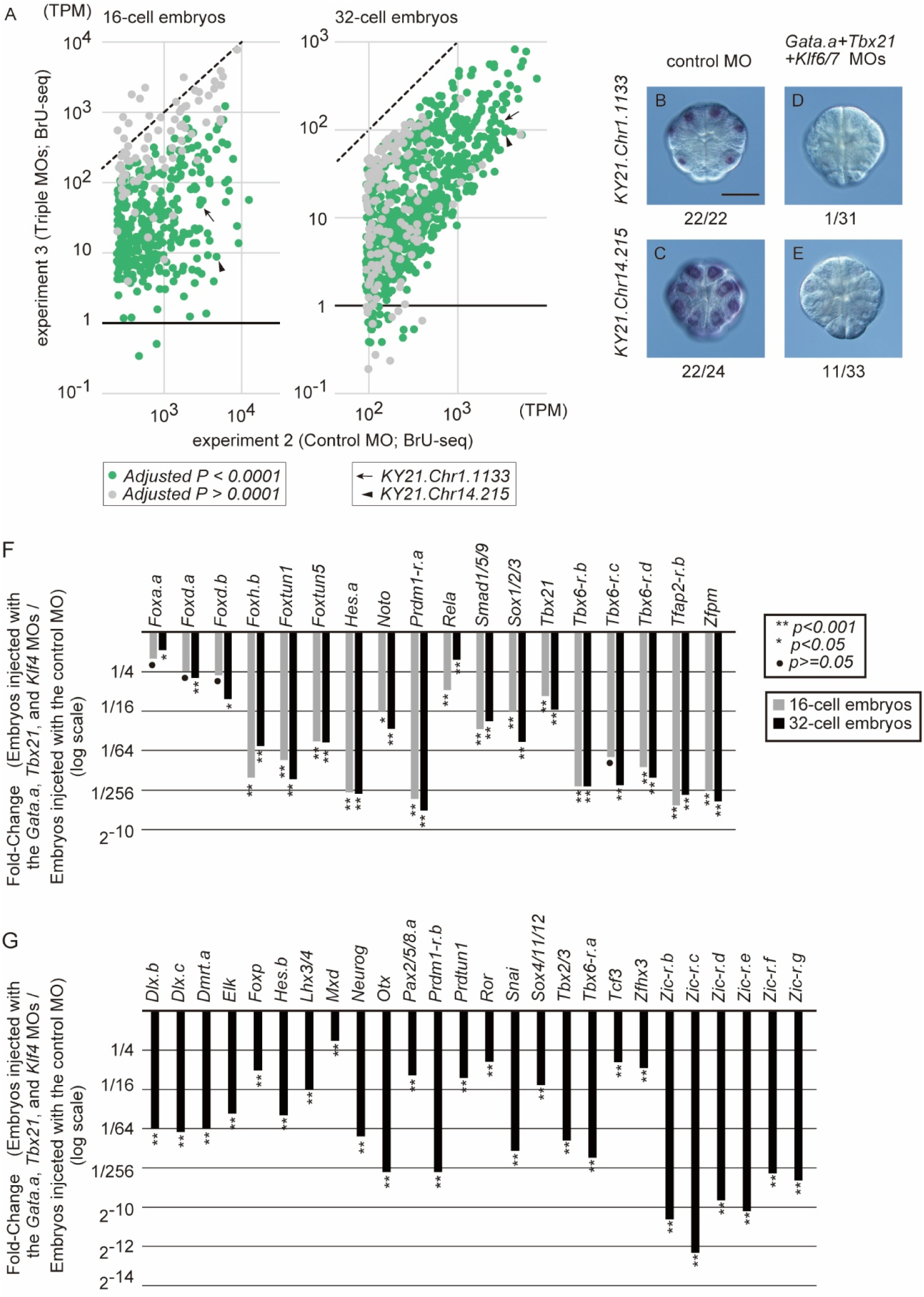
Genes downregulated in triple morphants of *Gata.a*, *Tbx21*, and *Klf6/7*. (A) Zygotically activated genes were plotted according to their TPM values obtained from experiments 2 and 3. Genes that were expressed differentially between control and triple-morphant embryos are shown by green dots and otherwise by gray dots. (B-E) Expression of two genes (KY21.Chr1.1133 and KY21.Chr14.215) in control and triple-morphant embryos was examined by *in situ* hybridization at the 16-cell stage. Total numbers of embryos and numbers of embryos with expression are shown beneath the photographs. Scale bar, 50 μm. (F) Changes in expression levels between control and triple morphant embryos were calculated using data obtained from the BrU-RNA-sequencing. Transcription factor genes that initiate expression from the zygotic genome at the 16-cell stage and at the 32-cell stage are shown. MOs were injected at 0.5 mM.

To validate the results of RNA-sequencing, we selected two genes (KY21.Chr1.1133 and KY21.Chr14.215) from the list of genes expressed at the 16-cell stage, and examined their expression using *in situ* hybridization. We chose these genes because their TPM values in experiment 2 were relatively high, and because their maternal expression was low according to previously published RNA-sequencing data for unfertilized eggs (Brozovic et al., 2018). While expression of these two genes was detected clearly in control embryos (Figure 5B, C), their expression was hardly revealed in triple morphant embryos of *Gata.a*, *Tbx21*, and *Klf6/7*, as expected (Figure 5D, E).

Because expression of transcription factor genes is well documented, we examined changes in expression levels of transcription factor genes in experiments 2 and 3 (Figure 5F, G). All transcription factor genes that initiate expression at the 16-cell stage were basically down-regulated in triple morphants at the 16-cell and 32-cell stages (Figure 5F, G). Exceptions are expression of *Foxa.a*, *Foxd.a*, *Foxd.b* and *Tbx6-r.c* in 16-cell embryos. These genes were indeed down-regulated, but statistically insignificantly.

Because the down-regulation of *Foxa.a* was the least pronounced, we examined its expression in 16-cell embryos by *in situ* hybridization. This gene is normally expressed in five pairs of cells called A5.1, A5.2, a5.3, a5.4, and B5.1 (Imai et al., 2004; Shimauchi et al., 1997), and this expression pattern was reproduced in embryos injected with the control MO (Figure 6A, B). In triple morphant embryos of *Gata.a*, *Tbx21*, and *Klf6/7,* this expression pattern was altered. Among 50 experimental embryos, 5 exhibited signals in A- and a-line cells, 41 only in A-line cells, and 4 gave no signals (Figure 6C, D). In addition, signals in these morphant embryos were clearly weaker than those in embryos injected with the control MO. Thus, it is likely that these three factors regulate *Foxa.a* expression differently according to cell lineages. This observation explains why the overall extent of *Foxa.a* down-regulation in the RNA-sequencing experiment was relatively modest.

**Figure 6.**
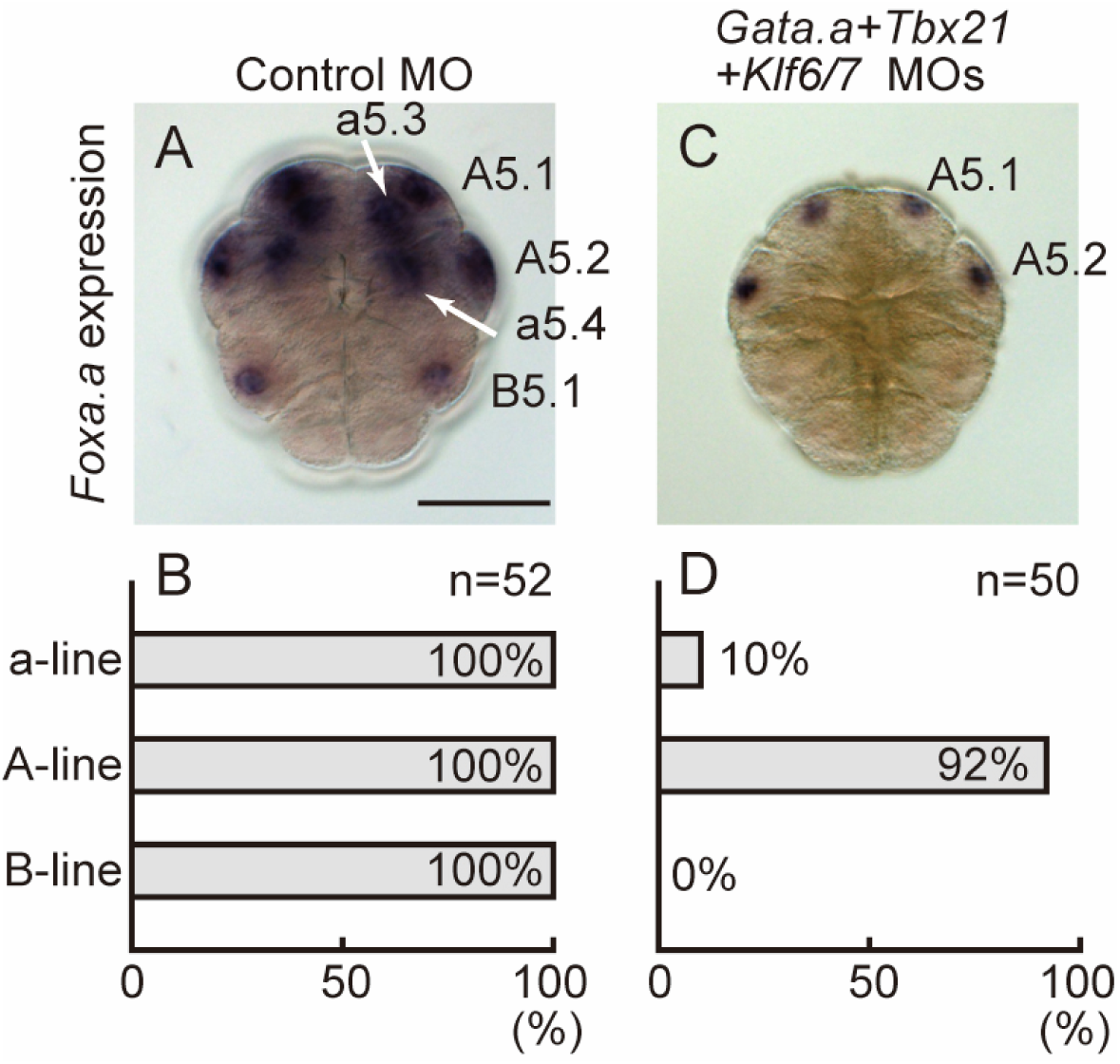
*Foxa.a* expression is abolished in a-line and B-line cells, but not in A-line cells of triple morphant embryos. (A, B) *Foxa.a* is expressed in A5.1, A5.2, a5.3, a5.4, and B5.1 cells of 16-cell embryos injected with the control MO (0.75 mM), and (C, D). It is expressed weakly in A5.1 and A5.2 cells, rarely in a-line cells, and it is absent in B-line cells of triple morphant embryos of *Gata.a*, *Tbx21*, or *Klf6/7* (0.25 mM each). (A, C) *Foxa.a* expression revealed by *in situ* hybridization. (B, D) Percentages of embryos that expressed *Foxa.a* in A-, a-, and B-line cells. Scale bar, 50 μm.

## Discussion

### Gata.a, Tbx21, and Klf6/7 are required for ZGA in ascidian embryos

Gata.a is distributed in all cells in early embryos (Oda-Ishii et al., 2016), and possesses at least two different functions (Imai et al., 2020). Specifically, it strongly activates target genes, including *Efna.d* and *Tfap2-r.b* in the animal hemisphere, and this activity is suppressed by β-catenin/Tcf7 in the vegetal hemisphere (Imai et al., 2020; Oda-Ishii et al., 2016; Rothbächer et al., 2007). At the same time, this factor provides basal activation of target genes even in the vegetal hemisphere, and contributes to their expression at physiologically normal levels (Imai et al., 2020). In the present study, we demonstrated that Tbx21 and Klf6/7 have functions similar to this secondary function of Gata.a. That is, Gata.a, Tbx21, and Klf6/7 are required for activating target genes at proper levels, and work cooperatively, although the relative contribution of each factor may vary depending on targets.

*Klf6/7* was not included in the lists of genes expressed zygotically in 16- or 32-cell embryos, suggesting that its regulatory role is mediated by maternally deposited mRNA and protein. Previous comprehensive assays have identified many maternal RNAs localized in the posterior vegetal region, and no other localization patterns have been reported (Ilsley et al., 2020; Matsuoka et al., 2013; Miwata et al., 2006). Because localization of *Klf6/7* is not reported in these studies, it is likely that maternal *Klf6/7* mRNA is distributed ubiquitously. Ubiquitous expression of maternal *Tbx21* (Takatori et al., 2004) and zygotic expression in all cells except germ-line cells (Imai et al., 2020) have also been reported. Therefore, it is likely that these two factors control levels of expression, but do not provide spatial cues.

On the other hand, *Foxa.a* expression in triple morphants was lost more frequently in B5.1 and a-line cells than in A-line cells. It is unlikely that Gata.a, Tbx21, and Klf6/7 confer spatial cues, because mRNAs for these factors are present in all blastomeres at almost the same levels. This suggests that *Foxa.a* expression in 16-cell embryos is regulated by multiple enhancers that regulate blastomere-specific expression, and that these enhancers differ in requirements for the three maternal factors.

The observation that 20% of zygotically activated genes were not under control of *Gata.a*, *Tbx21*, or *Klf6/7* indicates that there are enhancers that act independently of these three factors, as in the case of the *Foxa.a* enhancer that drives expression in A-line cells. It should be determined in future whether these 20% of genes require maternal factors other than Gata.a, Tbx21, and Klf6/7.

Gata.a, Tbx21, and Klf6/7 may be required for “priming” early enhancers before the 16-cell stage. Because Gata.a is present in nuclei as early as the 2-cell stage (Oda-Ishii et al., 2018), it likely binds to regulatory regions before ZGA, which occurs between the 8- and 16-cell stages. Because mRNAs for *Tbx21* and *Klf6/7* are also expressed maternally, Tbx21 and Klf6/7 may have such functions. This hypothesis is reminiscent of Zelda in *Drosophila* and Nanog, Pou5f1, and SoxB1 in vertebrate embryos (Gentsch et al., 2019; Lee et al., 2013; Leichsenring et al., 2013; Liang et al., 2008). These are all established pioneer transcription factors, and Gata.a and Klf4, which share a structural similarity with Klf6 and Klf7, are also pioneer transcription factors (Iwafuchi-Doi and Zaret, 2014). While it is unclear whether Tbx21 has such an activity, our results are consistent with the hypothesis that these three factors cooperatively create a transcriptionally competent chromatin state for approximately 80% of genes in 16- and 32-cell embryos.

### Highly-sensitive identification of zygotically activated genes in early embryos

Identifying zygotically activated genes in a comprehensive way is challenging, especially for zygotically activated genes that are expressed maternally at high levels. To overcome this, we previously utilized RNA-sequencing reads mapped to introns for detecting nascent transcripts (Imai et al., 2020). However, this assay had two limitations. First, because intronic transcripts rapidly degrade, even massive RNA-sequencing data may fail to identify relatively rare transcripts. Second, expression of single-exon genes or genes with short introns cannot be detected. In the present study, we used BrU-labeling to overcome these limitations.

For instance, while a gene encoding type-II cadherin was previously detected only at the 32-cell stage (Imai et al., 2020), zygotic expression of this gene was demonstrated at the 16-cell stage, consistent with *in situ* hybridization data (Imai, 2003; Noda and Satoh, 2008). In addition, the present study successfully identified zygotic expression of single-exon genes, including *Foxa.a*.

Despite its high sensitivity, this assay may still fail to identify certain zygotic genes, particularly those that are expressed maternally at extremely high levels, which will create intolerably high background signals that obscure low-level zygotic expression. Indeed, KY21.Chr7.1089 (KH.C7.190) was shown to be expressed zygotically in our previous study (Imai et al., 2020), but not in the present study. Nevertheless, by using 383 and 837 genes that we identified as zygotically activated at the 16- and 32-cell stages, respectively, we have shown that Gata.a, Tbx21, and Klf6/7 are required for proper expression of approximately 80% of genes at ZGA.

## Materials and Methods

### Animals and gene identifiers

Adult specimens of *Ciona intestinalis* (type A; also called *Ciona robusta*) were obtained from the National BioResource Project for *Ciona* (Satou et al., 2026). cDNA clones were obtained from our EST clone collection (Satou et al., 2005). Gene identifiers in the KY21 gene model set are listed in Table S9.

### Functional assays

The *Gata.a* MO was used in previous studies (Bertrand et al., 2003; Imai et al., 2016; Oda-Ishii et al., 2016; Rothbächer et al., 2007; Tokuoka and Satou, 2023). Sequences of other MOs, which block translation, are as follows: the primary MO for *Tbx21*, 5’-GATACTGGACTTCCGAAAAAGCCAT-3’; the primary MO for Klf6/7, 5’-ATATCCATGTTGTATGTTGGCACTC-3’; the secondary MO for *Tbx21*, 5’- AATAACTTAGAACCTTGTGCACCGT-3’; the secondary MO for *Klf6/7*, 5’-GTGGAAGTCACTCAAGTTACTACCT-3’. These MOs were microinjected into unfertilized eggs under a microscope. Concentrations of MOs are provided in the main text and/or figure legends. Whole-mount *in situ* hybridization was performed, as described previously (Satou et al., 1995). All functional assays were repeated at least three times using different batches of embryos.

### RNA-sequencing with BrU-injected embryos

BrU-labelled RNA molecules were obtained using the RiboCluster Profiler BRIC kit, (Medical & biological laboratories, #RN1008). Bromouridine was microinjected into unfertilized eggs at 100 mM. Immunoprecipitation of BrU-labelled RNA molecules was performed according to manufacturers’ instructions. Immunoprecipitated RNA fractions from two different conditions (experiments 2 and 3) and total RNA from uninjected control embryos (experiment 1) were converted to cDNA libraries using NEBNext Ultra II for DNA Library Prep kit (NEB, #E7645S).

Illumina sequencing reads were mapped to the KY21 gene model sets of the HT assembly of *Ciona robusta* (Satou et al., 2019; Satou et al., 2022) using Bowtie2 (Langmead and Salzberg, 2012). Differential gene expression was calculated with DESeq2 (Love et al., 2014). All *Ciona* proteins were compared with the human Uniprot proteome using BLASTP (UniProt-Consortium, 2019) (downloaded in March, 2021), and best-hit proteins with E-values less than 1×10^-5^ were regarded as homologs. GO annotations for human proteins were downloaded on March, 2021, from the Gene Ontology website (https://geneontology.org/) (Ashburner et al., 2000). Chi-square tests were performed to evaluate statistical significance.

RNA-sequencing data are deposited in the DRA/SRA database under accession numbers, DRR1076107–118.

## Acknowledgments

We thank Chikako Imaizumi (Kyoto University), Manabu Yoshida (University of Tokyo), and other members working under the National BioResource Project for *Ciona* (MEXT, Japan) at Kyoto University and the University of Tokyo for providing experimental animals. This research was supported by grants from the Japan Society for the Promotion of Science (24K02032, 24K21274 to Y.S, and 18H02376 to K.S.I.), and from Takeda Science Foundation to YS.

---

**Figure S1.**
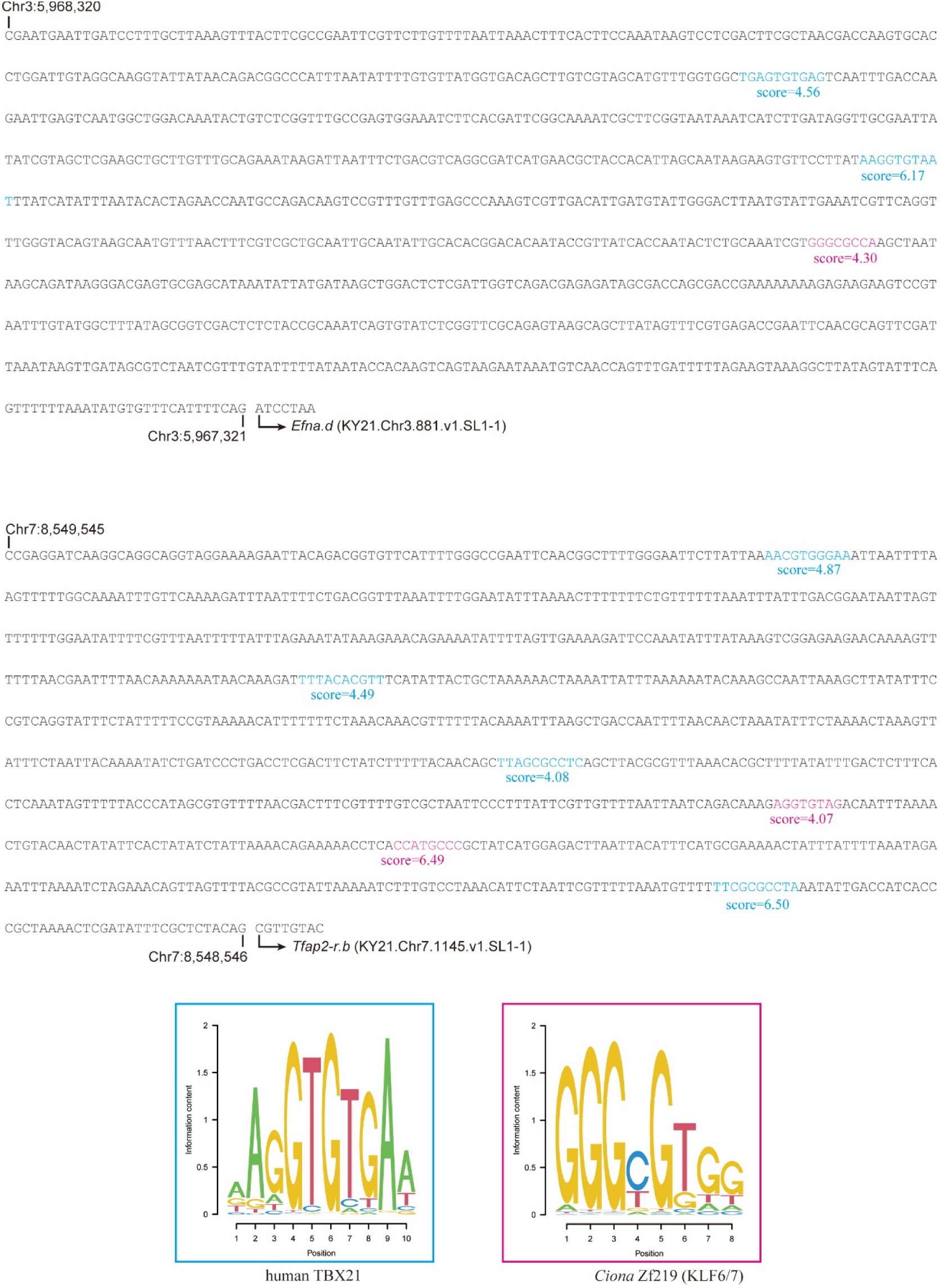
Upstream 1-kb regions of *Efna.d* and *Tfap2-r.b*. Putative Tbx21-binidng sites and Klf6/7-binding sites are shown by cyan and magenta letters with PWM scores calculated with the Patser program (Hertz and Stormo, 1999). Only sites with score >4 are shown. Position frequency matrices for human Tbx21 (JASPAR accession number MA0690.1) (Jolma et al., 2013; Sandelin et al., 2004) and for *Ciona* Klf6/7 (Nitta et al., 2019; Vincentelli et al., 2011) were used. These matrices were reproduced as sequence logos (Schneider and Stephens, 1990).

**Figure S2.**
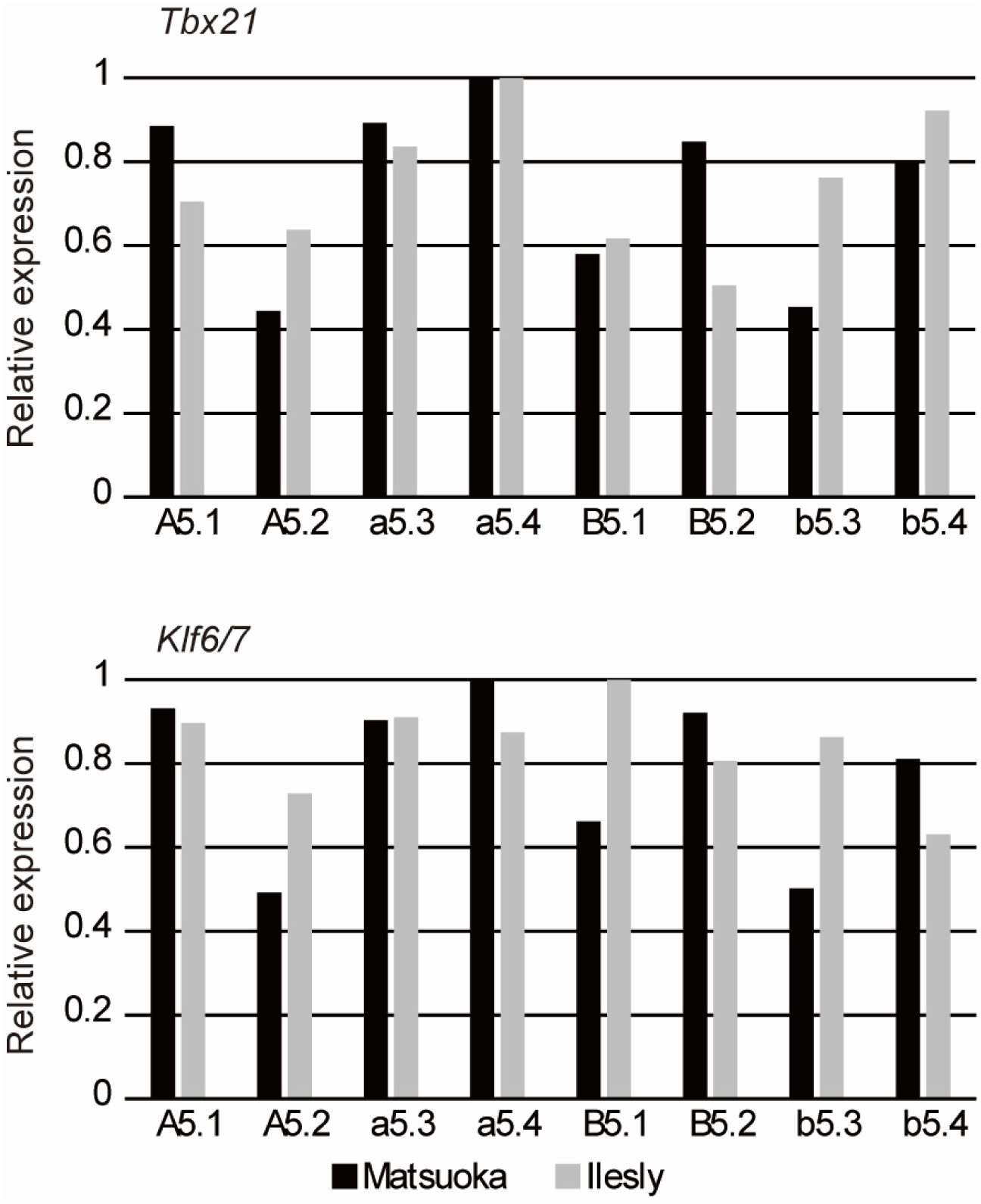
Expression levels of *Tbx21* and *Klf6/7* in 16-cell embryos. Relative expression levels of *Tbx21* and *Klf6/7* were calculated from microarray data (Matsuoka et al., 2013) and RNA-seq data (Ilsley et al., 2020), which measured mRNA expression levels in individual blastomeres, against the highest values in each dataset.

## Notes

### Competing Interest Statement

The authors have declared no competing interest.

